# Integrating patient and whole genome sequencing data to provide insights into the epidemiology of seasonal influenza A(H3N2) viruses

**DOI:** 10.1101/121434

**Authors:** Emily J. Goldstein, William T. Harvey, Gavin S. Wilkie, Samantha J. Shepherd, Alasdair R. MacLean, Pablo R. Murcia, Rory N. Gunson

## Abstract

Genetic surveillance of seasonal influenza is largely focused upon sequencing of the haemagglutinin gene. Consequently, our understanding of the contribution of the remaining seven gene segments to the evolution and epidemiological dynamics of seasonal influenza is relatively limited. The increased availability of next generation sequencing technologies allows rapid and economic whole genome sequencing (WGS). Here, 150 influenza A(H3N2) positive clinical specimens with linked epidemiological data, from the 2014/15 season in Scotland, were sequenced directly using both Sanger sequencing of the HA1 region and WGS using the Illumina MiSeq platform. Sequences generated by both methods were highly consistent and WGS provided on average >90% whole genome coverage. As reported in other European countries during 2014/15, all strains belonged to genetic group 3C, with subgroup 3C.2a predominating. Inter-subgroup reassortants were identified (9%), including three 3C.3 viruses descended from a single reassortment event, which had persisted in the population. Significant phylogenetic associations with cases of severe acute respiratory illness observed herein warrant further investigation. Severe cases were also more likely to be associated with reassortant viruses (odds ratio: 4.4 (1.3-15.5)) and occur later in the season. These results suggest that increased levels of WGS, linked to clinical and epidemiological data, could improve influenza surveillance.

## Introduction

Influenza viruses are a major cause of human morbidity and mortality worldwide, causing an estimated 250,000-500,000 deaths each year [1]. Influenza A virus (IAV) is an RNA virus, consisting of eight gene segments (PB2, PB1, PA, HA, NP, NA, M and NS). Classification of IAVs into subtypes is based on the combination of haemagglutinin (HA) and neuraminidase (NA) they possess. Influenza A viruses evolve rapidly by both mutation and reassortment. Mutations conferring incremental selective advantages result in the characteristic rapid antigenic drift of IAVs, while the segmented nature of the genome allows genomic reassortment when a cell is coinfected by two or more strains. Inter-subtype reassortment occasionally gives rise to viruses with pandemic potential [2], the most recent of which emerged in 2009 [3]. Intra-subtype reassortment is likely to occur much more frequently and play an important role in increasing viral genetic diversity and adaptive potential [4].

Surveillance is required to ensure that vaccine components reflect the antigenic characteristics of circulating IAV strains [2]. The HA is the primary antigenic determinant and consequently the chief focus of genetic surveillance, with influenza viruses routinely characterised into genetic groups based on amino acid residues of the HA protein, as defined by the European Centre for Disease Prevention and Control (ECDC) [5]. Consequently there is relatively little sequence data available for the remaining seven segments.

The West of Scotland Specialist Virology Centre (WoSSVC) is the Scottish influenza reference laboratory and is responsible for characterising several hundred IAV positive clinical specimens each year. Current characterisation of IAVs by the WoSSVC is based on Sanger sequencing of the HA1 region of the HA gene. Next generation sequencing (NGS) technology allows whole genome sequencing (WGS) of IAVs in a single reaction, permitting rapid and economical sequencing with the potential for high-throughput. In addition to viral characterisation, WGS enables the detection of reassortment events and antiviral resistance mutations anywhere in the genome.

The aim of this study was to assess the benefits of WGS over current Sanger sequencing methods for IAV surveillance and whether WGS can provide a greater understanding of the evolutionary and epidemiological dynamics of seasonal influenza. To this end, 150 influenza A(H3N2) positive clinical specimens from the 2014/15 influenza season in Scotland were sequenced using both methods. Genetic data and linked patient data were analysed to investigate rates of reassortment and potential associations between disease severity and phylogenies of each segment, reassortment status, and other patient details including location and age.

## Methods

### Samples

All 150 samples were influenza A(H3N2) positive clinical specimens submitted to the WoSSVC for routine influenza characterisation. Inclusion criteria for the study were all samples collected between 1st August 2014 and 31st May 2015 which had previously been genetically characterised using Sanger sequencing, providing enough material was available. Samples were received from 11 Health Boards throughout Scotland and included throat swabs (n=85), combined nose and throat swabs (n=15), gargle (n=14), nasopharyngeal aspirate (n=11), sputum (n=6), nasal swabs (n=5), tracheal aspirate (n=2), bronchoalveolar lavage (n=1) and non-classified respiratory specimens (n=11). The samples selected for Sanger sequencing at WoSSVC included a selection of sentinel surveillance samples collected from general practice surgeries (n=16), samples from patients with severe acute respiratory illness (SARI), as defined by Health Protection Scotland (n=22) and 112 other clinical cases.

### RNA isolation

Nucleic acid was extracted directly from clinical samples using automated extraction methods. Swabs, gargles and nasopharyngeal aspirates were extracted using the BioRobot MDx (Qiagen, Hilden, Germany) and sputum, tracheal aspirates and bronchoalveolar lavage were extracted using the NucliSENS easyMAG (bioMérieux, Marcy-l’Étoile, France). The same aliquot of extracted RNA was used for both Sanger sequencing and NGS.

### Sequencing of influenza viruses

Sanger sequencing was performed as previously described [6]. To prepare samples for NGS, RNA was reverse transcribed and the entire genome of influenza was amplified in a single RT-PCR reaction using the Uni/Inf primer set, as described by Zhou et al. [7]. Amplification was performed in 50 µl reactions containing 10 µl sterile water, 25 µl 2X RT-PCR buffer, 0.8 µl Uni12/Inf1 (10 µM), 1.2 µl Uni12/Inf3 (10 µM), 2 µl Uni13/Inf1 (10 µM), 1 µl SuperScript III Platinum Taq High Fidelity DNA Polymerase (Invitrogen, Carlsbad, CA, USA) and 10 µl RNA. Thermocycling conditions were as follows: 42°C for 60 min, 94°C for 2 min; 5 cycles (94°C for 30 s, 44°C for 30 s and 68°C for 3 min) followed by 31 cycles (94°C for 30 s, 57°C for 30 s and 68°C for 3 min), with a final extension step at 68°C for 5 min. DNA was diluted to a concentration of 175 ng in a volume of 50 µl and sheared acoustically using a Covaris S220 sonicator (Covaris, Woburn, MA, USA). NGS was performed on the Illumina MiSeq (Illumina, San Diego, CA, USA) as previously described by Wilkie et al. [8], with the following modifications: DNA was purified using 0.9 volumes of AMPure XP beads, adapter-ligated DNA was amplified using six PCR cycles and libraries were sequenced as 150-bp paired end reads.

### Bioinformatics

Illumina adapter sequences were removed from the data, paired-end reads were trimmed using a Phred score of 30 and to a minimum length of 50 bp using Trim Galore!(http://www.bioinformatics.babraham.ac.uk/projects/trim_galore/). Filtered reads were mapped to individual segments using the reference sequence A/Switzerland/9715293/2013. Two reference mapping software packages were employed: Tanoti (http://www.bioinformatics.cvr.ac.uk/tanoti.php) and Bowtie2 (http://bowtie-bio.sourceforge.net/bowtie2/index.shtml), default settings were used for both packages. Files were converted to BAM format using SAMtools (http://samtools.sourceforge.net/) and consensus sequences were obtained using DiversiUtils (http://josephhughes.github.io/DiversiTools/). Genome coverage and mean depth were greater using Tanoti, hence all further analyses were performed using consensus sequences obtained using this tool.

### Phylogenetic analysis

Consensus nucleotide sequences were aligned using MUSCLE [9]. Viruses were characterised into A(H3N2) genetic groups (subdivisions/subgroups) based on amino acid residues of the HA protein [5]. Time resolved phylogenetic trees were reconstructed using BEAST v1.8.2 [10]. The general time reversible model with a proportion of invariant sites and a gamma distribution describing among-site rate variation with four categories estimated from the data (GTR + I + Γ_4_) was identified as the best model of nucleotide substitution through comparison of Bayes factors [11]. Bayes factor analysis also determined that a relaxed (uncorrelated) molecular clock model [12] with branch rates drawn from a lognormal distribution, and a minimally constrained Bayesian skyline demographic model [13] should be used. Chains were run until convergence as identified using Tracer v1.6.0 and, after removing 10% of trees as burn-in, a sample of posterior trees was analysed using TreeAnnotator v1.8.2 to identify the maximum clade credibility (MCC) tree. The support for each node in the MCC tree is reflected by an associated posterior probability. Phylogenetic trees were visualised using the ggtree R package [14].

The positions of viruses of each genetic group were compared on phylogenies of each segment. Inconsistent positioning of single viruses or groups of viruses on these phylogenies was used to identify inter-subgroup reassortants. Phylogenetic mapping of reassortants was also performed computationally using the Graph-incompatibility-based Reassortment Finder (GiRaF) software v1.02 [15]. Posterior samples of phylogenies generated for each segment using BEAST were thinned to 1000 trees and analysed to identify both inter- and intra-subgroup reassortment events. Though every reassortment event must in reality split all eight segments into two subsets (retained and acquired segments), for some reassortment events the phylogenetic signal may be too weak for some segments to be assigned to one of the two reassorting sets of segments. Following the advice of the authors of the GiRaF software, to be considered a reassortment at least four segments were required to be confidently placed into either group.

Bayesian Tip-association Significance (BaTS) analysis [16] was used to detect significant phylogeny-trait correlations, testing the null hypothesis that there is no correlation between phylogeny and viral trait. Each gene segment was tested for a phylogenetic association with severity of infection (classified into SARI and non-SARI), patient age (categorised into <1 year, 1-5 years, 6-15 years, 16-64 years and ≥65 years), location by Health Board and genetic subgroup. The association index (AI) was calculated for each tree in a posterior sample generated by BEAST (after excluding the first 10% of tree states as burn-in) and compared to a null distribution generated by random reassignment of traits to tips of the phylogeny performed 5000 times. The AI ratio was calculated by dividing the observed AI by the null AI.

### Logistic Regression

Predictors of severe infection (defined by classification of a SARI case) were investigated by logistic regression using R v3.3.2 [17]. Sentinel surveillance samples were excluded from this analysis; the remaining 134 samples included 22 SARI cases. Patient age, location by Health Board, the week of sampling, genetic subgroup and whether the virus was classified as an inter-subgroup reassortant were tested as explanatory variables. To identify the best combination of explanatory variables, models were compared using AIC and likelihood ratio tests.

## Results

### Next generation sequencing of influenza A(H3N2) directly from clinical specimens

The mean coverage across the entire influenza A(H3N2) genome was 91%. Of the 150 samples sequenced, complete genome coverage was achieved for 71 samples and genome coverage of >90% was generated for 100 samples. Complete segment coverage was achieved for the two smallest segments, non-structural (NS) and matrix (M) protein, in all 150 samples. Segment coverage generally declined as the size of the segment increased, however average segment coverage of ≥80% was achieved for all segments (Table 1).

**Table 1.** Nucleotide coverage of gene segments of influenza A(H3N2) using next generation sequencing. RNA polymerase subunits (PB2, PB1 and PA), haemagglutinin (HA), nucleoprotein (NP), neuraminidase (NA), matrix protein (M) and non-structural protein (NS) segments (n=150).

| Segment | Size (nucleotides) | Mean coverage | Number of samples with 100% coverage |
| --- | --- | --- | --- |
| PB2 | 2280 | 80% | 82 |
| PB1 | 2274 | 83% | 74 |
| PA | 2151 | 89% | 105 |
| HA | 1701 | 96% | 135 |
| NP | 1497 | 94% | 126 |
| NA | 1410 | 98% | 142 |
| M | 982 | 100% | 150 |
| NS | 838 | 100% | 150 |
| Whole genome | 13133 | 91% | 71 |

Using NGS, the mean coverage of the HA gene was 1646 nucleotides, compared to an average length of 551 nucleotides when the HA1 region was sequenced using the Sanger method. When sequences generated from Sanger and NGS were compared >93% of the viral sequences had ≤2 amino acid differences in HA1, demonstrating a good correlation in the sequences obtained using both methods.

### Influenza A(H3N2) characterisation, epidemiology and resistance using NGS data

Full segment coverage of HA was achieved for 90% of samples using NGS and sufficient sequence data was available to characterise all 150 samples into genetic groups according to ECDC guidelines [5]. All viruses belonged to genetic group 3C, of which 107 viruses (71%) fell into genetic subgroup 3C.2a; five (3%) into subdivision 3C.3; six (4%) into subgroup 3C.3a, and 32 (21%) within subgroup 3C.3b. Nodes defining these four genetic groups on the HA phylogeny were associated with posterior probabilities exceeding 0.99, indicating strong support for the phylogenetic distinctiveness of each (Figure 1). Viruses of each genetic group did not appear to cluster geographically or in time during the 10-month study period (Figure 2), indicating co-circulation of distinct A(H3N2) lineages throughout the 2014/15 influenza season.

**Fig. 1.**
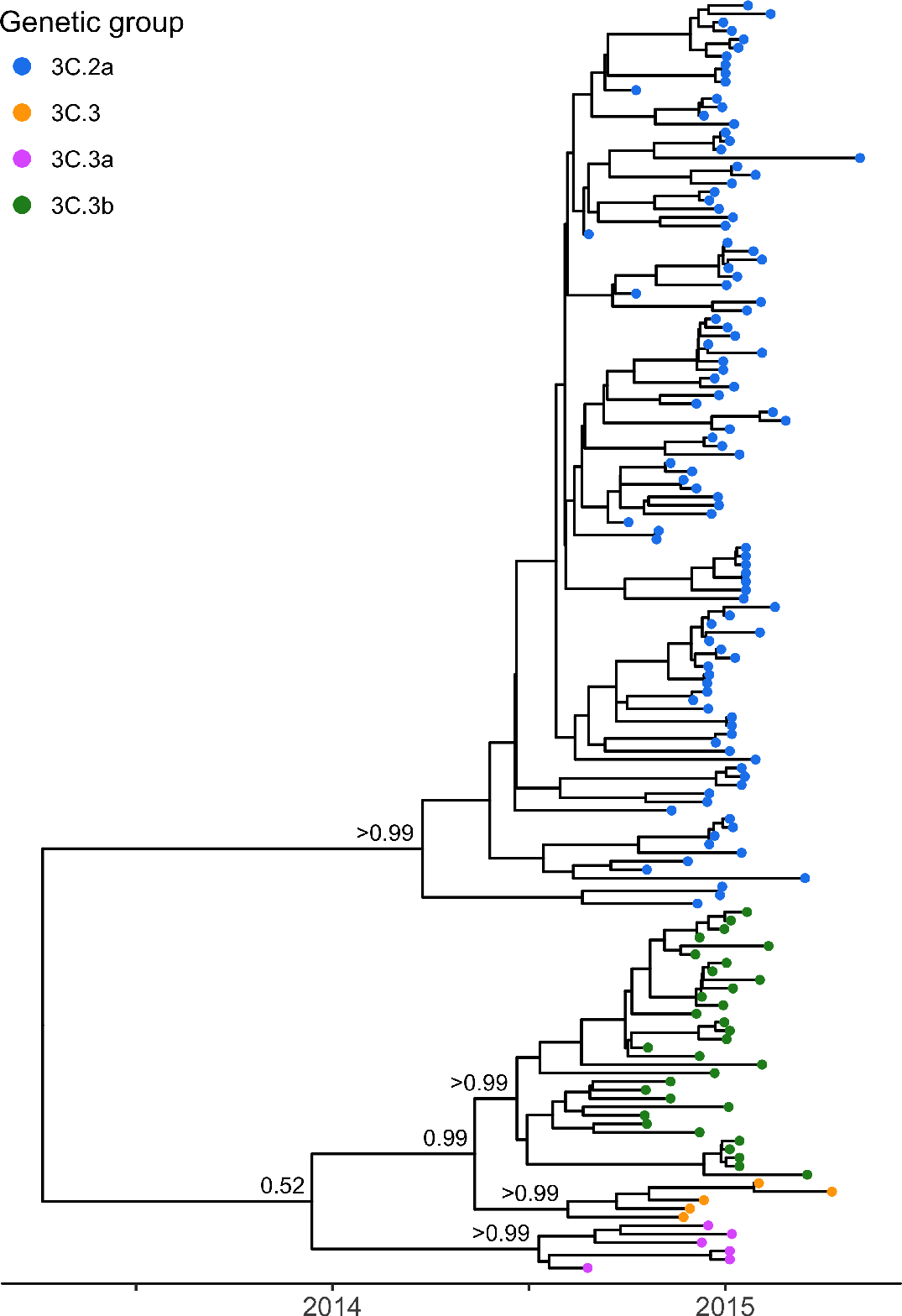
Phylogenetic tree of the haemagglutinin gene of 150 influenza A(H3N2) viruses obtained from the 2014/15 season. The maximum clade credibility, time-resolved phylogeny for consensus sequences of the HA gene obtained using NGS. Genetic group is indicated by colour. Posterior probabilities associated with the nodes defining each of the four genetic groups were >0.99.

**Fig. 2.**
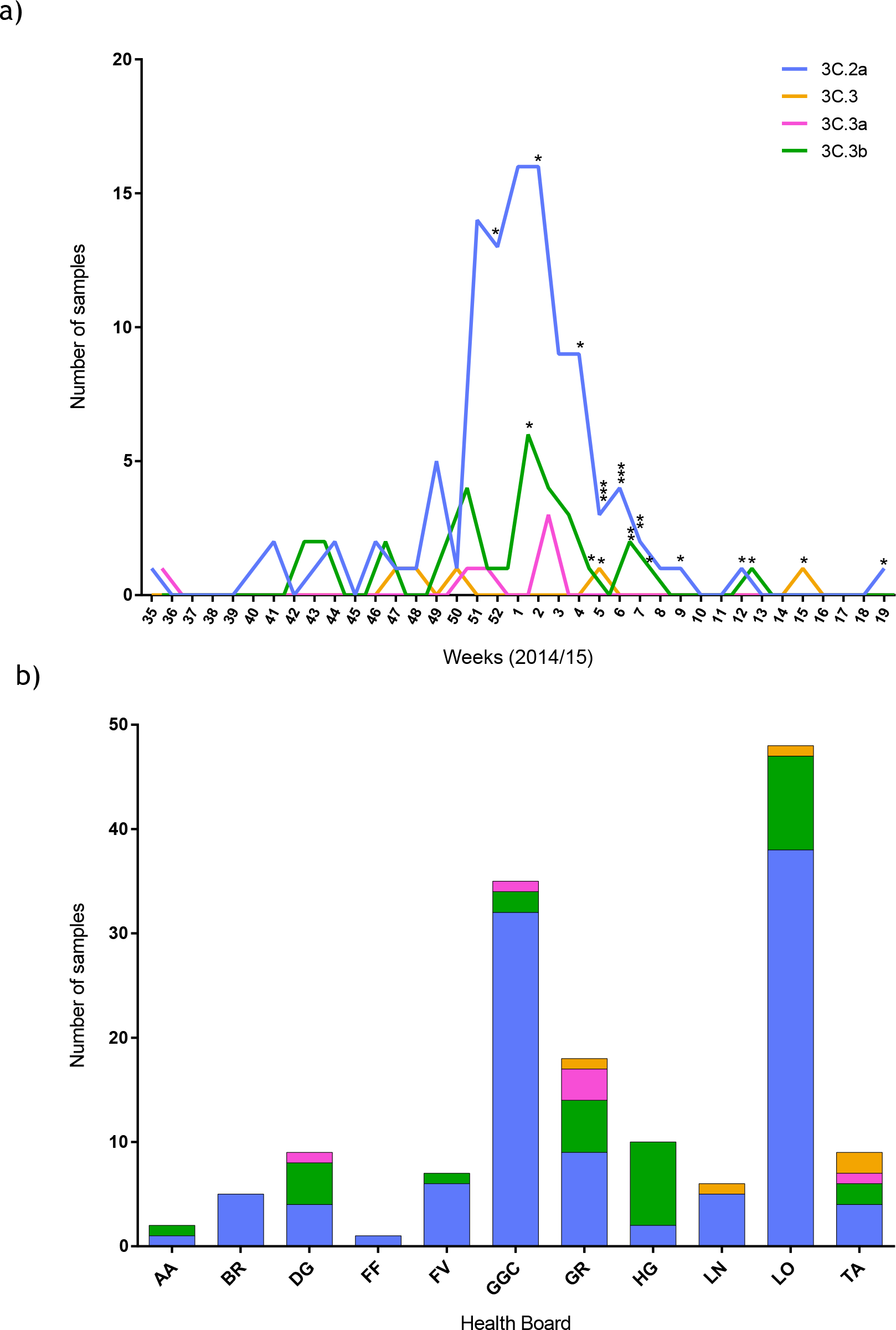
Circulation of 150 influenza A(H3N2) viruses throughout the 2014/15 influenza season. a) The number and genetic group of viruses received each week throughout the study period. Each asterisk represents a case of severe acute respiratory illness (SARI). b) Health Board from which the samples were collected. AA=Ayrshire and Arran, BR=Borders, DG=Dumfries and Galloway, FF=Fife, FV=Forth Valley, GGC=Greater Glasgow and Clyde, GR=Grampian, HG=Highlands, LN=Lanarkshire, LO=Lothian, TA=Tayside.

Whole genome sequencing enabled the presence or absence of drug resistance mutations in both the NA and the matrix 2 (M2) proteins to be determined. The S31N mutation in the M2 protein, which confers amantadine resistance, was present in all 150 samples, consistent with previous studies [18]. Substitutions in NA resulting in resistance to neuraminidase inhibitors; E119V, D151E, I222V, R224K, E276D, R292K, and R371K [19] were not detected (n=147).

### Analysis of whole genome sequence data and identification of reassortments

The sequences of all eight segments were concatenated to produce a single sequence for each virus and a whole genome phylogenetic tree was reconstructed. The concatenated genomes revealed a topology which was generally consistent with that of the phylogenetic tree generated using the HA gene only, with posterior probabilities of 1.00 on the nodes defining four distinct clades corresponding to genetic subgroups 3C.2a, 3C.3, 3C.3a and 3C.3b (Figure 3). The full genome phylogeny suggested the presence of some reassortant viruses and to determine the evolutionary relationships between gene segments, eight individual phylogenetic trees were generated (Figure 4). The position of viruses of each genetic group were compared on these phylogenies to identify inconsistencies arising from inter-subgroup reassortment.

**Fig. 3.**
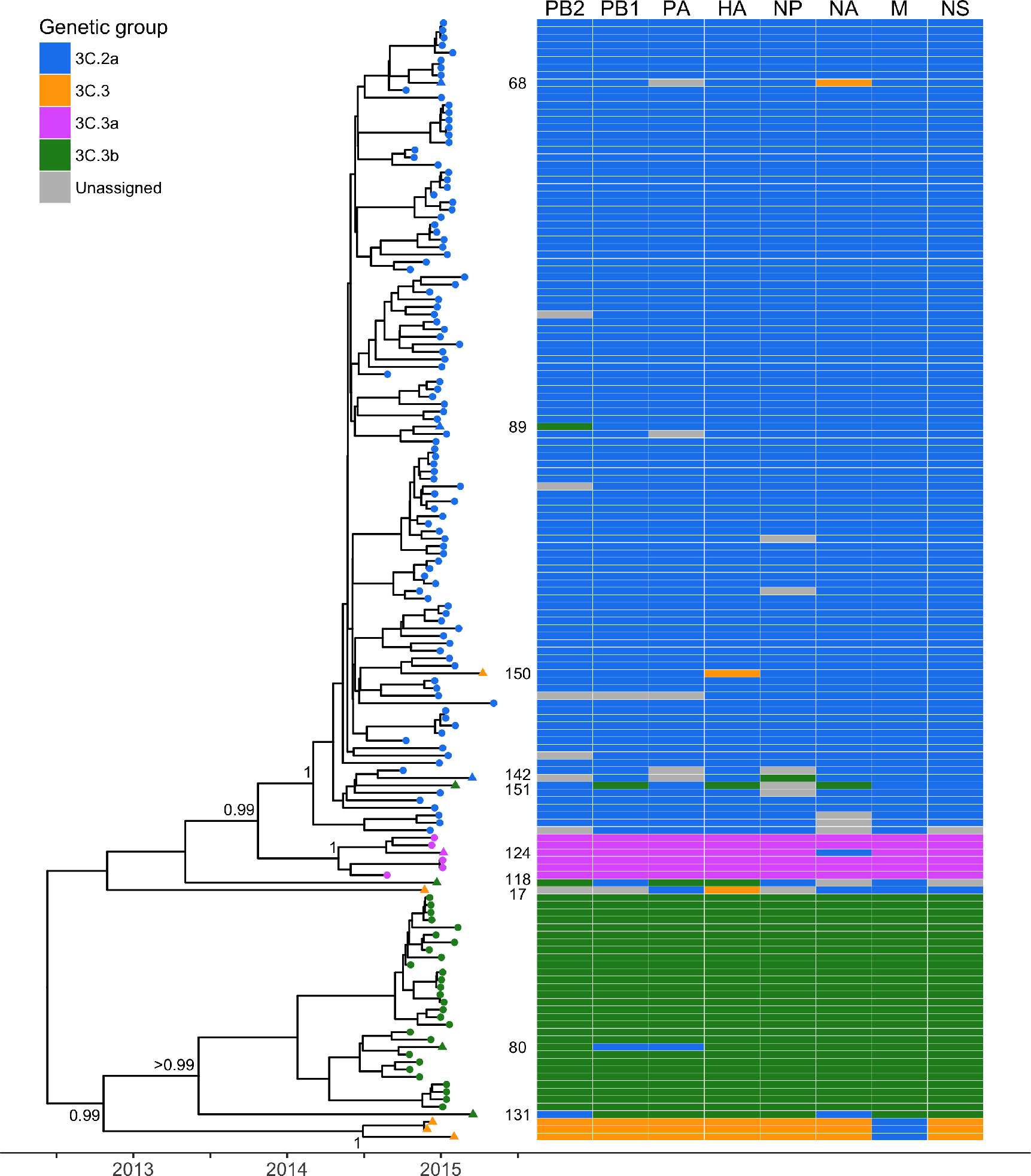
Phylogenetic tree of concatenated segments of 150 influenza A(H3N2) viruses from the 2014/15 season and schematic showing individual gene segment lineages. The maximum clade credibility, time-resolved phylogenetic tree was reconstructed for the whole genome of influenza A(H3N2) by concatenating all eight segments. Tips are coloured by genetic group (as characterised by HA sequence) according to the legend and triangles mark those identified as inter-subgroup reassortants (n=13). Sample numbers for these reassortants are also indicated, the three remaining unnumbered reassorted viruses make up the 3C.3 clade. Posterior probabilities associated with the nodes defining each of the four genetic groups were >0.99. To the right, a schematic representation of viral clustering of each gene segment is shown. Where samples could not be confidently assigned to a genetic group phylogenetically for a particular segment cells are coloured grey.

**Fig 4.**
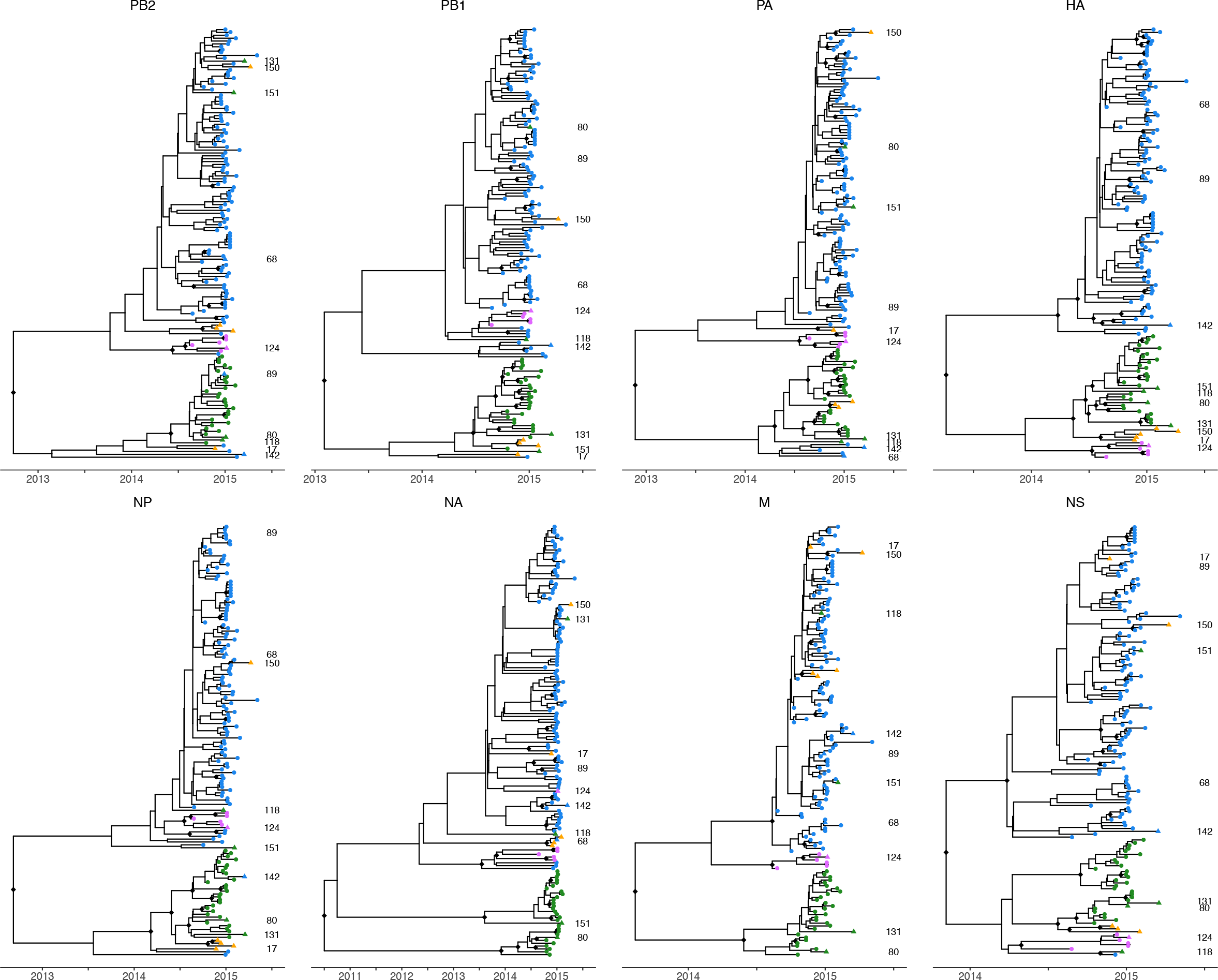
Phylogenetic trees of the eight individual gene segments from 150 influenza A(H3N2) viruses obtained from the 2014/15 season. The maximum clade credibility, time-resolved phylogenetic trees for consensus sequences of each segment obtained using NGS. Genetic groups are indicated on the tree by colour (3C.2a in blue, 3C.3 in orange, 3C.3a in pink and 3C.3b in green). The positions of novel inter-subgroup reassortant viruses in each phylogeny are indicated by triangles and are identifiable by sample number, the three remaining unnumbered reassorted viruses belong to the 3C.3 clade. Highly supported internal nodes of each phylogeny (posterior probability >0.9) are indicated by black diamonds.

Generally viruses belonging to the same genetic group, as characterised by HA sequence, also clustered together on phylogenies generated from each of the remaining seven gene segments. This suggests an absence of reassortment in the recent evolutionary history of the majority of viruses. However, 13 viruses were identified to be inter-subgroup reassortments (marked with triangles in Figures 3 and 4). A schematic representation of the position in the phylogenies of each of the eight gene segments is shown alongside the full genome tree in Figure 3. Ten of these reassortants were also detected using GiRaF, an automated reassortment detection tool. Of the remaining three (samples 17, 118 and 150), sample 118 was detected as a reassortant but below the significance threshold chosen. The GiRaF analysis additionally identified two intra-subgroup reassortment events in branches leading to two 3C.2a viruses and 18 3C.3b viruses respectively.

Ten of the 13 inter-subgroup reassortants were identified as descending from reassortment events leading to individual viruses in this study. The remaining three inter-subgroup reassortants, characterised as genetic subdivision 3C.3, were inferred to descend from a single reassortment event. These three viruses formed a distinct clade in each of the individual segment phylogenies, however the position of this clade in the overall topology varied. In seven of the eight segment trees these viruses were placed either outside the clades of other subgroups or within the 3C.3b clade, however in the M segment tree the 3C.3 viruses cluster within the 3C.2a viruses (posterior probability >0.99) (Figure 4). The M segments of these 3C.3 viruses were on average 10 and 14 nucleotides divergent from the M sequences of viruses belonging to genetic subgroup 3C.3a and 3C.3b respectively, and differed by only 4 nucleotides on average to the M segment of 3C.2a viruses. The three clinical samples harbouring these viruses were collected between November 2014 and April 2015, demonstrating that this reassortant genotype has persisted in the population. To compare these 3C.3 viruses detected in Scotland with those observed in the rest of the United Kingdom, all UK whole genome sequences from the 2014/15 influenza season available on GISAID (www.gisaid.org) were genetically characterised (n=171). Six of these viruses belonged to genetic subdivision 3C.3. Consistent with the pattern observed in the Scottish sequences, the M segment of these six viruses clustered amongst viruses characterised as genetic subgroup 3C.2a (data not shown).

### Analysis of virus phylogeny and trait correlations

To investigate predictors of a severe outcome in IAV infection, a logistic regression analysis was performed to analyse association between SARI cases and patient age, location, week of sampling, genetic subgroup and whether the virus was classified as an inter-subgroup reassortant (n=13). Week of sampling was found to be significantly correlated with severity of infection (likelihood ratio test (LRT), χ^2^=62.2, df=1, p<1×10^−10^), with SARI cases more likely to occur later in the season (as seen in Figure 2a). Severe cases were also found to be more likely to occur when patients were infected by a reassortant virus (LRT, χ^2^=4.9, df=1, p<0.05). The odds ratio for the association between inter-subgroup reassortants and SARI cases was calculated as 4.4 (95% CI: 1.3-15.5), or 4.8 (1.2-19.4) when only GiRaF-confirmed reassortments were considered (n=10). However, model selection indicated that the best model included week of sampling only; including reassortment status in addition to week did not result in an improved model (LRT, χ^2^=0.7, df=1, p>0.4). Further investigation showed that week of sampling was also correlated with reassortment status (LRT, χ^2^=7.3, df=1, p<0.01) therefore these could be confounding variables. No other explanatory variables tested were found to be significant (Table S1).

BaTS analysis was used to test for a phylogenetic association with severity of infection, patient age, location, and genetic group. Severity of infection was found to be strongly associated with the nucleoprotein (NP) phylogeny (AI ratio=0.76, p<0.01), and slightly weaker correlations with the M, HA, and NA phylogenies were also identified (p<0.05). Patient age was also found to be more clustered on the NP gene phylogeny than expected by chance (AI ratio=0.90, p<0.05). Location and genetic group were strongly associated with phylogeny for all eight gene segments (p<1×10^−10^). These correlations indicate strong geographic clustering not apparent at the resolution of genetic subgroup (Figure 2b) and that, as expected, viruses of the same genetic group (characterised by HA) tend to also have more similar sequences in other segments. BaTS analysis results are shown in full in Table S2.

## Discussion

The increased availability, decreased costs and turn-around times, both for sequencing and data analysis, of NGS is revolutionising microbiology. While Sanger sequencing of the HA1 region remains the predominant method used for IAV characterisation globally, this study demonstrates the applicability of NGS technology for influenza surveillance, allowing WGS directly from clinical specimens. The benefits of WGS over existing Sanger sequencing protocols for IAV surveillance include: 1. Greater resolution for genetic characterisation of IAV; 2. The level of drug resistance mutations in the NA and M segments can be evaluated; 3. Reassortment events can be detected and analysed; and 4. Mutations in any region of the genome not yet understood to be important (e.g. virulence factors) are available for retrospective analysis. Such retrospective analysis of mutations in any region of the genome other than HA1 is not available using current surveillance methods.

We demonstrate effective WGS direct from clinical specimens using only one nucleic acid extraction, RT-PCR and NGS reaction. Many previous studies have propagated patient isolates in cell culture prior to sequence analysis [20], however the results presented herein suggest that this additional step is unnecessary. In addition to the reduced time and costs involved, direct sequencing methods allow for analysis of non-culturable strains and avoid unwanted mutations that have been shown to occur during viral propagation [20-22].

Using NGS, complete coverage of the HA gene was achieved for 90% of samples and was adequate to allow characterisation of 100% of samples into genetic groups. When sequence data from Sanger and NGS were compared, high amino acid sequence homology was observed, providing further confidence that NGS could replace Sanger sequencing for routine influenza surveillance. Since 2016, ECDC guidelines have included HA2 residues for genetic characterisation of both influenza A(H3N2) and A(H1N1)pdm09 [5]. This means additional Sanger sequencing reactions are required, increasing time and costs, whereas these HA2 residues are routinely available with WGS, strengthening the case for NGS further.

The data presented reveal spatiotemporal co-circulation of distinct viral lineages, supporting previous data suggesting co-circulation of different A(H3N2) subgroups is common during epidemics of seasonal influenza [23-25]. This co-circulation facilitated inter-subgroup reassortment, estimated to occur at a rate of around 9% among the viruses studied. This supports previous data suggesting multiple reassortment events during an influenza season [25]. The level of inter-subgroup reassortment has been suggested to be an underestimate of true reassortment, as there may be undetected reassortment of segments between highly homogenous viruses of individual subgroups [26]. This was demonstrated here by the detection of additional intra-subgroup reassortments using computational detection methods. The 3C.3 lineage also demonstrates that inter-subgroup reassortment events can persist in the population and spread geographically. A total of nine viruses characterised as genetic subdivision 3C.3 (excluding novel reassortments) were observed (three and six from the Scottish and UK datasets respectively) in 2014/15. In all of these viruses the M segment clustered with viruses from genetic subgroup 3C.2a.

Persistence of intra-subtype reassortants has been demonstrated previously [27]. The factors associated with such persistence at a population level requires further investigation, however intra-subtype reassortment has been shown to temporarily raise the amino acid substitution rates contributing to an increased adaptive potential [28]. Specific examples of adaptive intra-subtype reassortment include a reassortment event between two antigenically distinct A(H3N2) lineages in 2003 that caused a major change in antigenic phenotype reducing vaccine effectiveness [23], and reassortment within A(H3N2) has also led to the global rise and spread of resistance to adamantane drugs [29]. Increased WGS over consecutive influenza seasons would allow for an increased understanding of the frequency and timing of such intra-subtype reassortment and the contribution to the evolutionary dynamics of seasonal influenza.

Logistic regression analysis indicated that infection with an inter-subgroup reassortant may be a risk factor for a severe outcome. It is possible that disruptions to inter-gene co-adaptations, caused by reassortment, could result in deviations from normal replication rates and virulence levels. With more data from WGS attached to patient information, this association could be investigated further. Both inter-subgroup reassortants and SARI cases were found to be more likely to occur later in the influenza season. It is possible that these correlations could result from a bias away from sampling milder cases later in the season. However, the distinct tendency for severe cases to occur later suggests that increased surveillance later in the season may be required to better understand the risk factors associated with disease severity.

There is currently limited data in the literature regarding risk factors associated with disease severity [30]. Broberg et al. [31] recently recommended influenza sequence data to be reported along with epidemiological data to allow for greater definition of factors which may increase the risk of severe influenza. BaTS analysis identified a significant association between phylogenies generated from the NP, M, HA, and NA gene segments and severe disease, with a particularly strong signal for NP. While these results should be interpreted with caution, they demonstrate the potential power of WGS coupled with linked epidemiological data. With more data, these correlations could be explored further to identify particular mutations in these genes which may be related to virulence.

In summary, this study demonstrates the benefit of NGS technology to provide whole genome sequence data for surveillance of seasonal influenza viruses. The results of both the logistic regression and BaTS analysis emphasise that WGS coupled to linked patient data could be an important tool for developing our understanding of the relationship between the influenza genome and disease severity. More generally, WGS provides the opportunity to further investigate the epidemiological consequences of within-subtype reassortment and both the intra- and inter-season evolutionary dynamics of seasonal IAV at the whole genome level.

## Acknowledgments

The authors would like to thank the Viral Genomics and Bioinformatics team at the MRC-University of Glasgow Centre for Virus Research for their assistance with bioinformatic analysis. We would also like to acknowledge Public Health England for uploading their full genome sequences to GISAID. Funding was provided by the MRC-University of Glasgow Centre for Virus Research and NHS Greater Glasgow and Clyde.

## Supplementary Materials

**Table S1.** Logistic regression model quality as assessed by AIC. Models are shown alongside values of AIC. Smaller AIC values indicate models of better quality (* marks best model). In models with a single explanatory variable, week is by far the most informative explanatory variable. Including reassortant or age in addition to week did not improve the model. Models including genetic subgroup or Health Board in addition to week could not be evaluated as fitted probabilities of 0 or 1 occurred, violating model assumptions.

| Model | AIC |
| --- | --- |
| Null intercept model | 121.7 |
| Severity ~ Health Board | 128.1 |
| Severity ~ Genetic subgroup | 123.7 |
| Severity ~ Age | 123.0 |
| Severity ~ Reassortant | 118.8 |
| Severity ~ Week* | 61.5 |
| Severity ~ Week + Reassortant | 62.8 |
| Severity ~ Week + Age | 63.4 |
| Severity ~ Week + Reassortant + Age | 64.7 |

**Table S2.** Summary of results of BaTS analysis.

| Trait | Segment | Association index (AI) ratio<sup>1</sup> | p-value<sup>2</sup> |
| --- | --- | --- | --- |
| Severity of infection | PB2 | 0.96 | 0.37 |
|  | PB1 | 0.94 | 0.30 |
|  | PA | 0.92 | 0.22 |
|  | HA | 0.80 | 0.03 |
|  | NP | 0.76 | <0.01 |
|  | NA | 0.80 | 0.03 |
|  | M | 0.82 | 0.01 |
|  | NS | 1.00 | 0.50 |
| Age | PB2 | 0.92 | 0.07 |
|  | PB1 | 0.92 | 0.08 |
|  | PA | 0.93 | 0.08 |
|  | HA | 0.93 | 0.11 |
|  | NP | 0.90 | 0.02 |
|  | NA | 0.93 | 0.12 |
|  | M | 0.95 | 0.11 |
|  | NS | 0.94 | 0.09 |
| Health Board | PB2 | 0.62 | <1x10<sup>-10</sup> |
|  | PB1 | 0.63 | <1x10<sup>-10</sup> |
|  | PA | 0.71 | <1x10<sup>-10</sup> |
|  | HA | 0.62 | <1x10<sup>-10</sup> |
|  | NP | 0.70 | <1x10<sup>-10</sup> |
|  | NA | 0.62 | <1x10<sup>-10</sup> |
|  | M | 0.78 | <1x10<sup>-10</sup> |
|  | NS | 0.69 | <1x10<sup>-10</sup> |
| Genetic subgroup | PB2 | 0.16 | <1x10<sup>-10</sup> |
|  | PB1 | 0.10 | <1x10<sup>-10</sup> |
|  | PA | 0.13 | <1x10<sup>-10</sup> |
|  | HA | 0.13 | <1x10<sup>-10</sup> |
|  | NP | 0.14 | <1x10<sup>-10</sup> |
|  | NA | 0.13 | <1x10<sup>-10</sup> |
|  | M | 0.11 | <1x10<sup>-10</sup> |
|  | NS | 0.13 | <1x10<sup>-10</sup> |

<sup>1</sup>. Observed AI/null AI where null AI is derived from 5000 tree tip randomisations. Lower values indicate stronger phylogeny-trait associations in observed data.
<sup>2</sup>. BaTS null hypothesis test.

